# Geometric constraint of mechanosensing in bone marrow stromal cell cultures prevents stiffness-induced differentiation

**DOI:** 10.1101/2023.10.05.560162

**Authors:** M Hernandez-Miranda, D Xu, DA Johnston, M Browne, RB Cook, BG Sengers, ND Evans

**Affiliations:** Centre for Human Development, Stem Cells and Regenerative Medicine, Bone and Joint Research Group, Institute for Life Sciences, University of Southampton Faculty of Medicine, Southampton; Bioengineering Sciences Research Group, University of Southampton Faculty of Engineering and Physical Sciences, Southampton; Biomedical Imaging Unit, University of Southampton Faculty of Medicine, Southampton

**Keywords:** Stiffness, bone marrow stromal cells, differentiation, elastic modulus, thickness, osteogenic, hydrogel

## Abstract

Extracellular matrix (ECM) stiffness is fundamental in cell division, movement and differentiation. The stiffness that cells sense is determined not only by the elastic modulus of the ECM material, but also by ECM geometry and cell density. We hypothesised that these factors would influence cell-traction-induced matrix deformations and cellular differentiation in bone marrow stromal cells (BMSCs). To achieve this, we cultivated BMSCs on polyacrylamide hydrogels that varied in elastic modulus and geometry and measured cell spreading, cell-imparted matrix-deformations and differentiation. At low cell density BMSCs spread to a greater extent on stiff compared to soft hydrogels, or on thin compared to thick hydrogels. Cell-imparted matrix deformations were greater on soft compared to stiff hydrogels or thick compared to thin hydrogels. There were no significant differences in osteogenic differentiation relative to hydrogel elastic modulus and thickness. However, increased cell density and/or prolonged culture significantly reduced matrix deformations on soft hydrogels to levels similar to those on stiff substrates. This suggests that at high cell densities cell traction-induced matrix displacements are reduced by both neighbouring cells and the constraint imposed by an underlying stiff support. This may explain observations of the lack of difference in osteogenic differentiation as a function of stiffness.

## INTRODUCTION

The ability of cells to sense and respond to mechanical information from the extracellular matrix (ECM) is important for many biological processes ^1^. ECM stiffness directs stem cell spreading, differentiation, migration, and proliferation ^2–4^ and is now one of the most studied ECM mechanical properties. ECM stiffness has been shown to be of fundamental importance in specifying stem cell differentiation ^5,6^ and so the mechanical properties of tissues likely play a fundamental role in tissue development and regeneration.

Adherent cells sense the stiffness of their growth substratum by applying traction forces at their points of attachment and by sensing their dynamic displacement as a function of the applied force. Generally, on stiff materials, the resistance to this applied force results in assembly of focal adhesions, F-actin and the generation of intracellular tension. This subsequently results in cell spreading. On soft materials, however, cells are unable to generate internal tension and as a result spread to a lesser degree ^2^. Downstream signalling then regulates gene expression and directs cell migration, proliferation, and differentiation in a process known as mechanotransduction ^7^.

The ECM stiffness that an adherent cell senses is not only dependent on the elastic modulus of the material, however. It may also be dependent on the dimensions of the material, the magnitude of the applied load and dynamic changes in material structure that may occur on timescales similar to those of cell induced tractions. For small cells on large material structures, this is often negligible. However, it becomes important when, for example, the thickness of a soft material adherent to a stiff underlying support is reduced. Many studies of cell mechanobiology use ECM-modified polyacrylamide (PAAm) hydrogel as a cell culture substrate, a material that displays largely linearly elastic behaviour. For cell studies, this material is polymerised *in situ* and bound to a glass or plastic support for ease of handling. Several studies have shown that for a hydrogel with a defined, low elastic modulus, the cell begins to behave as if it is on a much stiffer material as the thickness (or depth) of the material decreases ^8–10^. This is due to the constraint to cell-induced lateral hydrogel deformation imposed by the underlying stiff support – effectively the cell must induce a greater strain in a thin material than in a thick material for an identical *lateral* surface displacement ^11^. This also becomes important on thicker materials when cells begin to act collectively. Trepat *et al* noted that large colonies of MDCK cells were insensitive to substrate stiffness ^12^, and subsequently we found that this scaled with colony size ^13^, reflecting earlier work that demonstrated that groups of cells contract materials to a much greater degree than individual cells ^14–16^. In effect, colonies exert much larger lateral tractions than individual cells, and thus ‘feel more deeply’ into materials. In addition, cells are able to sense and respond to the dynamic disturbances neighbouring cells impart on a common ECM, and cells can become mechanically-coupled through the ECM material in the absence of any cell contact – cells can ‘feel each other’ through the material ^17^. As groups of cells exert more strain on their materials than individual cells, these collective groups can feel each other at greater distances than isolated, individual cells. Taken together these data indicate that collective behaviour of cells and material dimensions must be considered when interpreting mechanobiological observations.

The effect of ECM stiffness on bone-derived stem cells has been widely studied due to its importance in skeletal repair and in tissue interactions with orthopaedic biomaterials. Increasing stiffness is generally correlated with increased osteogenic differentiation in cell populations containing putative skeletal stem cells in the published literature (for example, bone marrow stromal cells (BMSCs) or mesenchymal stem cells (MSCs); reviewed in ^18^). However many of these studies examine differentiation at low cell density ^5,19,20^, and some studies conclude that matrix stiffness may not correlate with osteogenic differentiation, with some cell populations remaining insensitive ^21–26^. Recently, Venugopal *et al.* showed the importance of cell density in differentiating human MSCs ^27^. At lower cell densities, MSCs spread and divided significantly less on soft compared to stiff substrates. However, as cell density increased, cell spreading and division was increased on soft materials, and become similar regardless of the substrate stiffness. This indicates the likelihood that individual MSCs are able to sense the dynamic mechanical strains imparted on materials by neighbouring cells, which they interpret as an increase in stiffness. It remains unknown, however, how the ability of monolayers of cells to sense substrate stiffness may be impacted by changes in hydrogel dimensions.

In this study, we tested the hypothesis that reduced substrate thickness limits the ability of BMSCs to deform polyacrylamide substrates and reduces their ability to mechanosense soft materials and differentiate accordingly. To achieve this we modulated hydrogel thickness, elastic modulus and cell density and compared cell differentiation and cell-induced hydrogel displacements.

## METHODS

### Fabrication of polyacrylamide hydrogels

Polyacrylamide (PAAm) hydrogels were prepared according to the method of Pelham and Wang (Pelham and Wang, 1997). Glass coverslips (13 mm or 25 mm diameter) (VWR International, Leicestershire, UK) were used as rigid support for the hydrogels were cleaned with tissue paper and functionalised with 0.1 M NaOH (Sigma-Aldrich, Gillingham, UK), on a plate heater at 80 °C for 20 min. Next, coverslips were washed with distilled water, dried before covering the surface with 3-aminopropyltriethoxysilane (APES) at room temperature for 5 min (Sigma-Aldrich) and rinsed with distilled water. Dried coverslips were immersed for 30 minutes in 0.5% (v/v) glutaraldehyde (Sigma-Aldrich) in phosphate-buffered saline (PBS) (Sigma-Aldrich), washed and dried again. Hydrogels with different elastic moduli were prepared by varying the concentration of acrylamide-bisacrylamide. 12.5% (v/v) acrylamide, 1.5% (v/v) bisacrylamide and 85% (v/v) PBS for soft hydrogels and 20% (v/v) acrylamide, 24% (v/v) bisacrylamide and 55% (v/v) PBS for stiff hydrogels. The mixture was degassed for 15 minutes under a vacuum. 0.1% (v/v) of N, N, N’, N’-tetramethylethane-1,2-diamine (TEMED) and 1% (v/v) of solution of 10% (w/v) ammonium persulfate (APS) (Sigma-Aldrich) was added to the mixture and vortexed to initiate the polymerisation. Specific mixture volumes were situated between a pre-treated coverslip and glass. Once the hydrogels polymerised, they were immersed in PBS for 10 minutes, carefully separated from the glass slide, placed on well plates with PBS and washed overnight at 4 °C. Hydrogels were washed 3 times with new PBS, covered with sulfosuccinimidyl 6(4-azido-2-nitrophenyl-amino) hexanoate (sulfo-SANPAH) (ThermoFisher Scientific, Loughborough, UK) 0.5 mg/mL in 4-(2-hydroxyethyl)-1-piperazineethanesulfonic acid (HEPES) and exposed to UV light (Chromato-vue TM-20, UVP transilluminator, 240 V) for 25 minutes. Later, the hydrogels were washed 3 times with HEPES 50 mM pH 8.5, and 0.1 mg/mL collagen solution type I (Sigma Aldrich) was added to cover the hydrogels before incubating overnight at 4 °C. For hydrogels containing fiduciary markers, fluorospheres at a concentration of 1% (v/v) (ThermoFisher Scientific) of 0.5 µm diameter were included in the PAAm mixture before polymerisation and sonicated before use. For measurements of hydrogel thickness, allylamine (Sigma-Aldrich) was added at 0.196% v/v to the acrylamide-bis-acrylamide mixture before polymerisation ^13^. Once the PAAm hydrogels polymerised, hydrogels were incubated in Alexa Fluor 568 (Thermo Fisher Scientific) 1 mg/mL (1:50) at room temperature for 3 h before washing 3 times with PBS 1×.

### Measurements of hydrogel thickness

Hydrogel thickness was measured by confocal microscopy (Leica TCS SP5, Leica, Cambridge, UK). Soft and stiff PAAm hydrogels of different thicknesses (3 samples per condition) on 13 mm glass coverslips were placed upside-down on a thin glass slide and immersed in 1× PBS. Hydrogels were imaged at 20× magnification and 2µm or 10 µm *z*-stacks from top to bottom. The fluorescent intensity profiles were obtained and analysed using the Leica Software (LAS X Core Offline) to calculate the thickness (*z*-value). The images were used to quantify manually the number of wrinkles on the hydrogel surface.

### Measurements of hydrogel elastic modulus

Soft and stiff PAAm hydrogels with different thicknesses were fabricated as previously described, and stiffness was measured using a nanoindenter (NanoTest Vantage system; MicroMaterials Ltd., Wrexham) as described in Xu *et al*., 2023 ^28^. In brief, the samples immersed in PBS solution were tested using a spherical diamond tip (500 µm). Nanoindentation was carried out in load control to various maximum loads (10 μN to 850 μN, minimum load step: 2 μN) to obtain the indentation modulus (Er) vs depth (δ) profile. The indentation depth/hydrogel thickness ratios (δ/h) varied between 0.01 and 0.5, with the indents spaced apart by 250 µm. The maximum load was applied for 120 s at rates of 1 µN/s for loading and 5 µN/s for unloading at (20 ± 1 °C).

### Cell culture

The MG63 human osteosarcoma cell line (Sigma, St. Louis, USA) was obtained from the Health Security Agency, Porton Down, UK and used at passages <30. BMSCs were previously isolated from human bone marrow samples obtained from the Spire Southampton Hospital and the Southampton General Hospital under local ethical approval. As previously described ^29^, Stro-1^+^ BMSCs (passage 1-5) were grown in α-MEM media and MG63 cells (variable passage number) in DMEM with 10% v/v Fetal Bovine Serum (FBS) and 100 µg/mL penicillin/streptomycin on tissue culture polystyrene flasks and incubated at 37 °C. The medium was changed every 2 - 3 days.

### Actin, nuclei, and vinculin staining

Vinculin was stained to identify focal adhesions in Stro-1^+^ BMSCs by immunocytochemistry. Cells were fixed in 4% (w/v) paraformaldehyde (PFA) for 20 minutes at room temperature, rinsed with PBS 1X and permeabilised with 0.5 % (v/v) Triton, X-100 in PBS for 30 minutes at the same temperature. After washing 3 times with PBS 1×, cells were incubated in 0.1% (w/v) BSA in PBS at 4 °C for 2 hours. Next, cells were incubated with the primary vinculin rabbit anti-human, mouse, polyclonal antibody (ThermoFisher Scientific) at 2 µg/mL (final concentration) in 0.1% (w/v) BSA in PBS overnight at room temperature. Cells were washed with 0.1% (w/v) BSA in PBS 3 times and incubated with the goat anti-rabbit IgG (H+L) highly cross-adsorbed secondary antibody, Alexa Fluor Plus 594 at 10 µg/mL (final concentration) for 1 hour at room temperature. FITC-conjugated phalloidin (ThermoFisher Scientific) (1:1000) was added to stain actin fibres and incubated for 30 minutes at room temperature (foil covered) and finally rinsed 3 times with PBS 1X. Thirdly, cells were stained with DAPI for 5 min at room temperature to stain nuclei and finally rinsed with PBS twice. Cells were imaged using a Nikon Ti inverted microscope and Leica confocal microscope with filters for DAPI (em ʎ = 350 ± 50 nm; ex ʎ = 460 ± 50 nm); eGFP (em ʎ = 470 ± 40 nm; ex ʎ = 525 ± 50 nm); Cy3 (em ʎ = 545 ± 25 nm; ex ʎ = 605 ± 70 nm), and Cy5 (em ʎ = 620 ± 60 nm; ex ʎ = 700 ± 75 nm); filters and merged using ImageJ/FIJI v. 2.9.0 free software.

### Cell spreading area quantification

SSCs were plated on PAAm hydrogels and tissue culture polystyrene, incubated for 24 hours at 37 °C, and placed under a Nikon Ti inverted microscope. 5 microscopic fields of each soft and stiff hydrogel of different thicknesses (n=3) and phase-contrast images were imaged at 10× magnification. The pictures were analysed using the FIJI software by drawing around the cell periphery using the wand (tracing) tool and by using the analysis menu for quantification. The cell area was measured using the analysis menu. The number of cells varied depending on the hydrogel elastic modulus and thickness.

### Time-lapse imaging and digital image correlation

Time-lapse imaging was used for displacement microscopy studies to determine how cells perceive the rigidity of materials in different conditions, as described previously ^13^. Briefly, cells plated on soft and stiff PAAm hydrogels with different thicknesses were photographed every 5 minutes for 24h starting at different time points (days 1, 7, 10, 14, week 7, if applied) using a Nikon Eclipse Ti inverted microscope. The fluorescent images were acquired using the Cy3 channel (em ʎ = 545 ± 25 nm; ex ʎ = 605 ± 70 nm) and were saved as an ND2 file (Nikon, UK). The images were extracted from the ND2 file and uploaded into a MATLAB algorithm developed by Zarkoob et al ^14^ using digital image correlation to track displacements. Firstly, a grid (10 × 8) was drawn to yield 99 nodes for the analysis. The tracking parameters were Kernel size=31; Subpixel size=9; Smoothness=5; maxMove=6; smoothGrid= 25. MATLAB (The Math Works R2017a, Natick, MA) was then used to quantify the total cumulative displacements from the first image taken at 5 min (image 1) to the following taken every hour (image 2, 3, …, 24, etc.) in each node, by registering and adding up the changes in displacements between images taken one hour apart. The cumulative displacement data from the 99 nodes in 3 hydrogel triplicates were used to calculate the 90^th^ percentiles and standard deviation at each hourly time point. The data were plotted in GraphPad Prism v.10 as XY graphs, and the average of the 90^th^ percentile from ∼8 h to 24 h as grouped graphs.

### Osteogenic differentiation and alkaline phosphatase (ALP) activity quantification

Cells were plated at 5,000 cells/cm^2^ on TCP and PAAm hydrogels. 1 day post seeding ascorbate-2-phosphate, beta glycerophosphate and dexamethasone were added to basal medium final concentrations of 280 µM, 5 mM and 10 nM, respectively. Medium was changed every 2 or 3 days for 7 or 14 days, before ALP biochemical quantification. A basal medium was used as a negative control. For ALP staining, medium was removed from the cells on TCP or PAAm hydrogels on day 7 or 14 and washed twice with PBS. Ethanol (95% v/v) was added to each well and incubated for 10 minutes at 4°C. Ethanol was aspirated, and the plate was washed twice with PBS before drying at room temperature. Fast violet (Sigma Aldrich) [0.24 M] was dissolved in α-naftol solution in dH_2_O (4% v/v), added to the cells, and incubated for 1 hour at 37 °C before removing the solution. The wells were rinsed with 1 mL Milli-Q water and imaged using a Zeiss microscope with a colour camera included (Axiovision software). For ALP quantification, medium was removed from the wells before adding 500 µL cell Lytic (Sigma-Aldrich) and incubated for 15 minutes. Later, the suspension was recovered and centrifuged at 4 °C at 80 g for 15 minutes. Finally, the cell suspension was recovered and transferred to new tubes, frozen at -80 °C until their use. For the ALP quantification, the assay buffer for the standards and the ALP substrate solution (0.04 g phosphatase substrate (Sigma Aldrich) in 10 mL alkaline buffer solution (1.5 M) (Sigma Aldrich)). Later, p-nitrophenol standards at different concentrations were prepared. For the assay, 100 µL standards, 20 µL of cell lysate and 80 µL substrate were added in triplicates to a 96 transparent well plate. Additionally, 100 µL NaOH 1 M was added to each background control. After that, the well plate was incubated at 37 °C for 60 min or until samples acquired a yellow colour, and the reaction was finished with 100 µL NaOH. The absorbance was quantified in a Glomax reader at 405nm.

## RESULTS

### Control of polyacrylamide (PAAm) geometry and stiffness

To test the effect of hydrogel elastic modulus and thickness on the morphology, differentiation, and cell-generated tractions of BMSCs, we first prepared PAAm hydrogel substrates suitable for cell culture. The bulk modulus and thickness of substrates were controlled by varying the cross-linker and monomer concentration and volume solution, respectively, to produce ‘soft’ and ‘stiff’ hydrogels (of predicted elastic modulus 1 kPa and 40 kPa, respectively) ^2,13^. The thickness of both ‘soft’ and ‘stiff’ hydrogels increased as a function of volume of polymer reaction solution used to form the gels, from 54.5 ± 5.7 µm to 597 ± 15 µm for soft hydrogel and from 27.4 ± 1.6 µm to 277 ± 25 µm for stiff hydrogels (Figure 1A). The thickness of soft hydrogels was significantly greater than that of stiff hydrogels when made at a polymerisation solution volume of 50 µL but not at any other volume. However, there was a general trend for stiff hydrogels to be thinner than soft hydrogels. Close examination of hydrogels by confocal sectioning revealed the presence of surface wrinkles on soft hydrogels, which were absent on stiff hydrogels (Figure 1B). Wrinkles on thin materials were more tortuous and shorter than on thicker materials; in the latter, continuous wrinkles often extended over significant parts of the hydrogel surface. Reflecting this, the number of discrete wrinkles declined due to increasing thickness (Figure 1C). In parallel, we used nanoindentation to measure substrate stiffness including a correction for hydrogel thickness ^28^. For ‘soft’ hydrogels formed from polymerisation volumes ≥ 25 µL corresponding to thicknesses > 200 µm, there were no significant differences in measured stiffness, with mean values of 5.50 ± 2.0 kPa, 5.27 ± 1.5 kPa and 5.51 ± 0.17 kPa. However, for thin hydrogels made from 5 µL (∼50 µm), the modulus was slightly but significantly higher than for any other at 6.95 ± 0.15 kPa (p < 0.001). As expected, the reduced modulus values for ‘stiff’ hydrogels were much greater than those for soft hydrogels but with no significant difference between any group with values of 55.7 ± 0.89 kPa, 56.2 ± 0.42 kPa, 53.69 ± 1.41 kPa and 59.77 ± 0.74 kPa for gels made from 5, 25, 50 and 100 µL gel solution, respectively.

**Figure 1.**
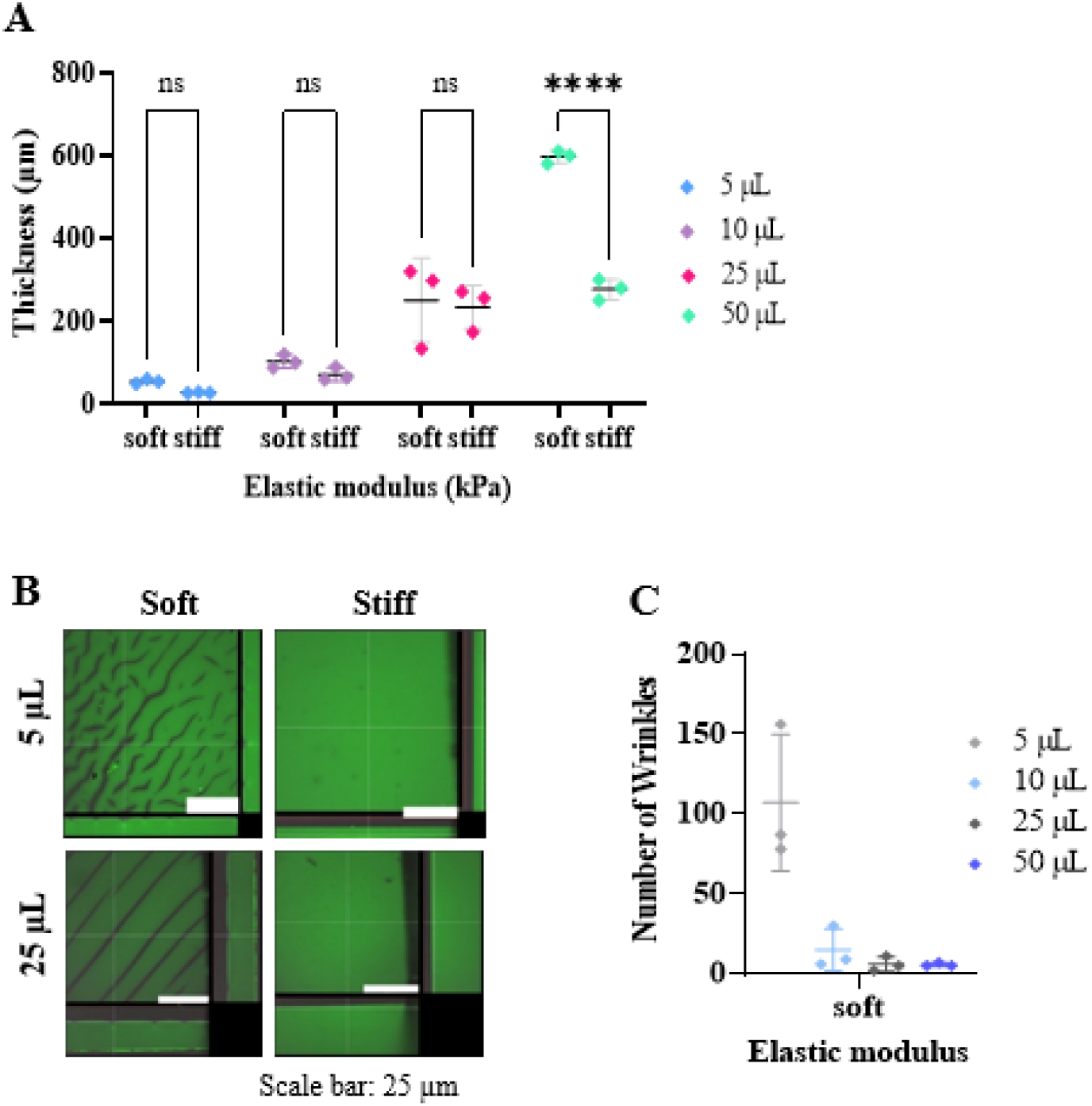
Hydrogel thickness is proportional to the polyacrylamide mixture volume and influences the hydrogel surface and elastic modulus. A) Polyacrylamide hydrogel thickness varies proportionally to the mixture volume, and soft hydrogels are thicker than their stiff counterparts at 50 µL. Thickness was measured by confocal microscopy. B) Soft polyacrylamide hydrogels exhibit surface wrinkles, whereas the surface of stiff polyacrylamide hydrogels remains intact. Large images show x-y optical sections at the surface of the PAAm hydrogel marked with a line in the bottom and side z-images C) Wrinkles on soft polyacrylamide hydrogels per field of view decrease relative to PAAm volume. A 2-way ANOVA was used to test for significant differences (n=3). ****= p<0.0001.

### Hydrogel elastic modulus and thickness modify the spreading cell area of BMSCs

In previous work by ourselves^13^ and others^5^, increasing stiffness leads to cell morphology and spreading changes. To confirm this, we plated primary BMSCs on the fabricated substrates and quantified cell morphology by microscopy. Cells stained with labelled phalloidin and immunostained for vinculin on stiff substrates ∼200 µm in thickness appeared larger in area than those on soft materials, with more prominent pseudopodia (Figure 2A). This was confirmed by quantifying cell area. Cells on stiff substrates had larger spreading areas (3230 ± 160 μm^2^) compared to cells on soft (1780 ± 1260 μm^2^) hydrogels (Figure 2B, p < 0.05). Previous studies by us^2^ and others^4^ found that substrate thickness dictates the apparent stiffness sensed by cells. To test this, we compared cell morphology and spreading on ‘soft’ hydrogels of thickness ∼50 µm compared to ∼200 µm. Cells on thin materials exhibited greater cell spreading area observed by microscopy (Figure 2C), an observation confirmed by quantification of cell area, where cells on thin hydrogels spread to a greater degree (4840 ± 3040 μm^2^) compared to cells on thick hydrogels (1780 ± 1260 μm^2^; Figure 2D). Although cells on stiff materials or soft, thin substrates had significantly larger areas, the data tended to group into two populations, reflecting heterogeneity in cell response.

**Figure 2.**
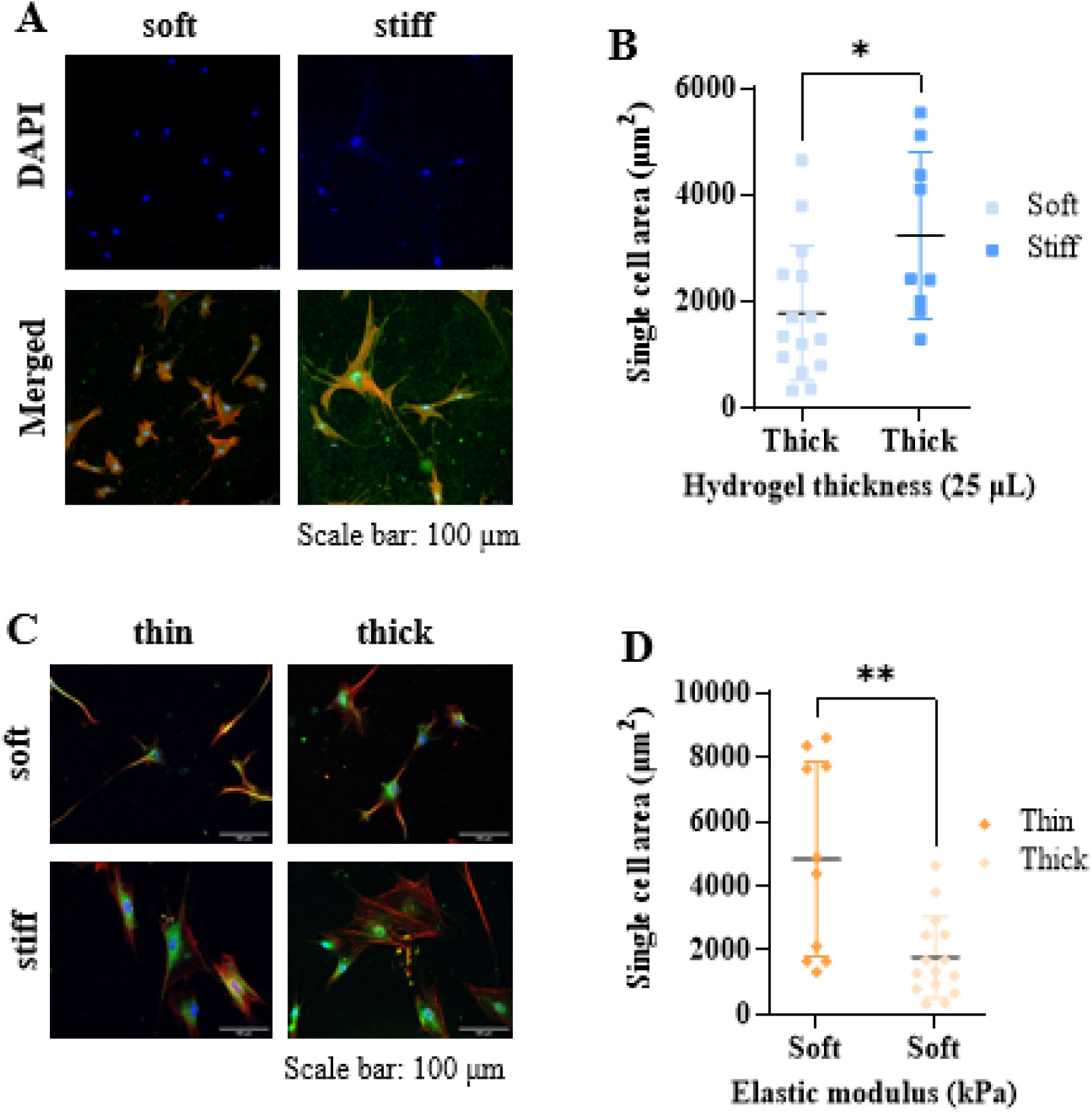
Bone marrow stromal cells mechanosense changes in elastic modulus and thickness and modify their spreading area and morphology in response. A) Nucleus (blue), actin (red) and vinculin (green) staining of BMSCs on PAAm hydrogels with different elastic moduli. Small cells are visible on soft hydrogels, whereas larger cells are seen on stiff hydrogels. B) BMSCs exhibit a greater cell spreading area on stiff compared to soft PAAm hydrogels (* < 0.05). C) Nucleus (blue), actin (red) and vinculin (green) staining of BMSCs on soft, stiff, thin, and thick PAAm hydrogels. Actin fibres were clear in cells on stiff materials. D) Cells spread more on soft, thin compared to soft, thick counterparts (** < 0.01). A Student’s t-test was used to calculate statistical significance of differences of means.

This data shows that MSCs respond to increased detected substrate stiffness by increased spreading, either because of intrinsic bulk substrate elastic modulus or reduced substrate thickness.

### Reduced substrate thickness restricts cell-induced substrate tractions

Increased cell spreading as a function of decreasing substrate thickness is likely due to the underlying glass surface’s constraint of cell-imparted hydrogel tractions. The cell interprets this as an increased material stiffness, despite the bulk modulus of the soft hydrogels remaining unchanged ^11^. To test this hypothesis, we compared cell-induced displacements at the hydrogel surface of thin and thick, soft, and stiff hydrogels by time-lapse displacement microscopy (note that this is an equivalent method to traction force microscopy, but in this study, we did not compute traction forces).

To achieve this, it was first necessary to include fluorescent particles in the hydrogels as fiduciary markers for digital image correlation analysis of time-lapse images (Supplementary Figure 1A). Including these particles did not alter the thickness of the PAAm hydrogels, except for a volume of 50 µL hydrogel solution (Supplementary Figure 1B). Fluorescent particles appeared to accumulate in areas of gel wrinkles and were observable as defined lines in fluorescent images of the gels. The addition of fluorescent particles caused minor changes in gel stiffness relative to controls, but did not affect the relative stiffness of ‘soft’ compared to ‘stiff’ hydrogels (Supplementary Figure 1C and D).

Next, we plated BMSCs on thin (5 µL, ∼50 µm) or thick (25 µL, ∼200 µm) soft hydrogels and quantified cell-induced displacements over 24 hours. Five minutes after cell seeding, BMSC morphology was similar regardless of the hydrogel’s elastic modulus or thickness (Figure 3A and phase contrast videos in Supplementary Video 1). However, it was evident from video microscopy of the fluorescence-particle labelled hydrogels that cells induced much larger tractions on thick substrates than those on thin substrates (Supplementary Video 1). Note that aggregations of fluorescent particles were evident in areas where cells were not present, as well as in areas where they were, the former reflecting pre-cell seeding hydrogel swelling and the latter reflecting cell-induced gel contraction occurring before the start of video microscopy. To test the hypothesis that cells induce larger tractions on thick compared to thin substrates quantitatively, we measured hydrogel displacements and compared them between substrates. Hydrogel displacements increased over time in all cases, rising more quickly and to higher values on soft vs stiff materials and thick vs thin materials (Figure 3B). When comparing cumulative displacements over the 24-time period, displacements were significantly greater for soft, thick substrates than soft, thin substrates (Figure 3C). Despite the very low observable deformations, this trend was also apparent for stiff substrates (images and videos for stiff materials are available in Supplementary Figure 2 and Supplementary Video 2).

**Figure 3.**
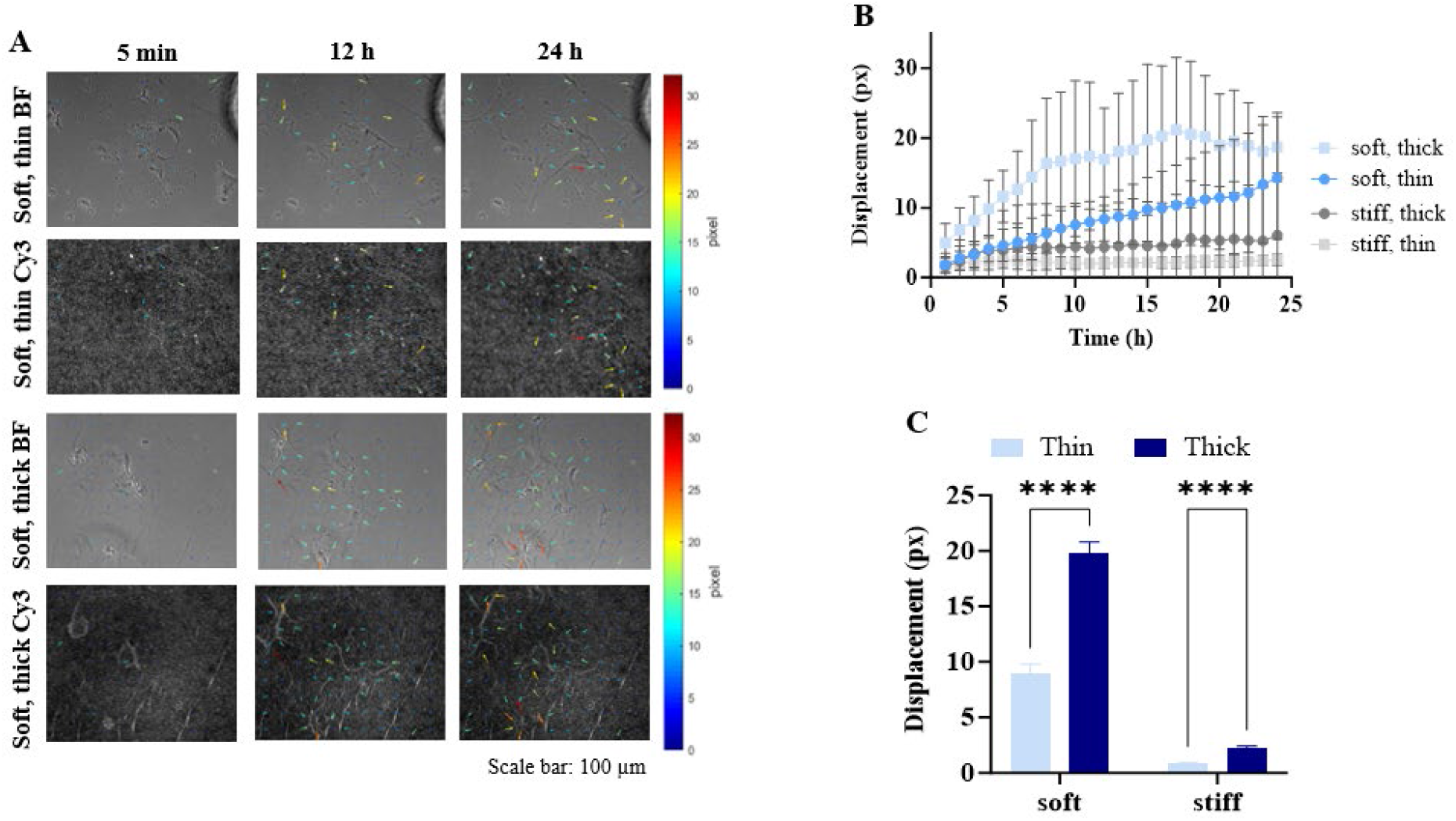
Hydrogel deformations by BMSCs increase over time and depend on the hydrogel’s mechanical properties. A) Time-lapse imaging of the Stro-1^+^ BMSCs on soft, thin and thick PAAm hydrogels with embedded fluorescent microbeads. Cell movements promote wrinkle formation over time; displacements indicated with coloured arrows. Note that arrow length was scaled by a factor of 3 for visualisation. Phase contrast (BF) and fluorescent (Cy3) images were obtained at 10X magnification under a Nikon Eclipse Ti inverted microscope. Scale arrow=3. B) Hydrogel displacements increase over time and are greater on soft, thick hydrogels than on stiff matrices. Squares represent the mean and SD of the 90th percentiles of displacements of the 99 nodes (n= 3 hydrogels). C) Stro-1^+^ BMSCs generated more significant deformations on thick than thin PAAm hydrogels, regardless of the hydrogel rigidity. A two-way ANOVA was used to assess statistical significance of difference of means; **** p< 0.0001.

These data confirm that the surface tractions induced in soft hydrogels by BMSCs are reduced by decreasing hydrogel thickness due to constraints imposed by the proximity of the underlying glass surface to which the hydrogel is attached.

### The osteogenic differentiation of BMSCs does not depend on material elasticity and thickness

BMSCs are known to differentiate into functionally distinct lineages due to changes in substrate elastic modulus, with increased differentiation into osteoblastic lineage cells on stiffer compared to softer materials ^5^. To test whether BMSCs would perceive a reduction in the thickness of a material with a low elastic modulus as an increase in stiffness, we plated BMSCs on thick or thin substrates of high and low elastic modulus and measured osteogenic differentiation. As expected, osteogenic supplements induced the differentiation of primary BMSCs to the osteogenic lineage, as measured by alkaline phosphatase activity, regardless of substrate elasticity. However, in contrast to our hypothesis, we found no significant differences in differentiation because of increased bulk elastic modulus or because of decreased substrate thickness (Figure 4).

**Figure 4.**
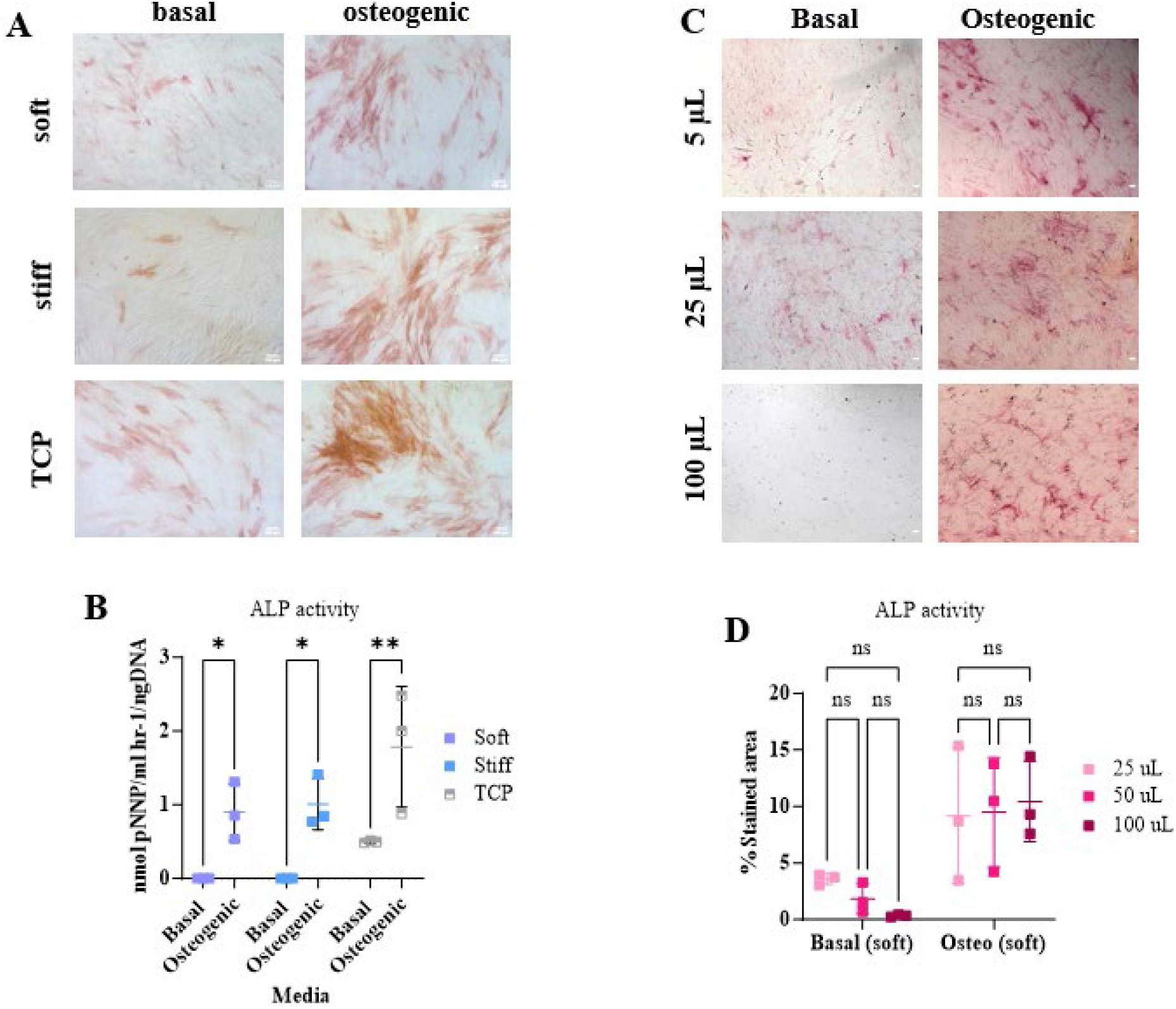
Substrate stiffness and thickness do not increase the osteogenic differentiation of BMSCs. A) Osteogenic supplements but not substrate stiffness enhances the ALP activity of MSCs, shown as red/brown staining. B) ALP activity increased under osteogenic conditions compared to basal media; non-significant changes were detected with increasing stiffness. Squares represent the mean ALP activity normalised to DNA content and standard deviation (****=p<0.001). C) ALP staining increased in osteogenic compared to basal medium, but no clear differences as a function of thickness were seen D) The addition of osteogenic supplements but not the change in substrate thickness increases the ALP activity of cells. Squares represent the mean and standard deviation of the percentage of the stained area. The 2-way ANOVA method was used to calculate significant differences.

### Cell-induced traction forces decrease with respect to cell density

One explanation for the lack of differentiation in marrow stromal cells as a function of stiffness or thickness may be constraints imposed by neighbouring cells at high density. As reported in a previous study ^27^, this effect has been hypothesised to reduce the ability of cells to mechanosense soft materials due to competing tractions from neighbouring cells ^17^. In addition, large cumulative tractions induced on substrates by monolayers of cells may result in insensitivity to substrate elastic modulus due to whole-scale contraction of the hydrogel by collective cell action. ^13^ To directly test this, we plated BMSCs at increasing densities on soft, thick materials (283 ± 50 µm) and quantified hydrogel displacements at 20 hours. As predicted by Venugopal et al.^27^, displacements declined as a function of cell density (Figure 5A and B and Supplementary Video 3). Cells at 1,000 cells/cm^2^ created greater hydrogel displacements (21.8 ± 1.1 µm) compared to the displacements generated by cells at 20,000 cells/cm^2^ (6.3 ± 0.4 µm).

**Figure 5.**
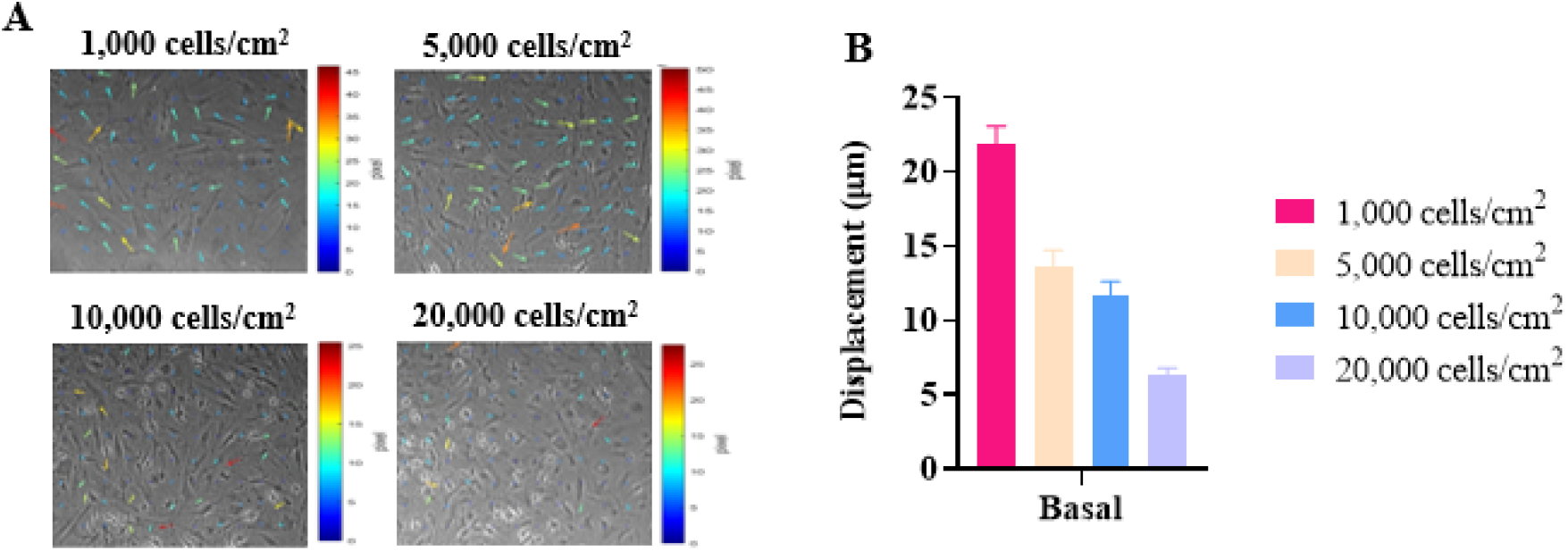
Matrix deformations decrease with respect to increased BMSCs seeding density. A) Time-lapse imaging of MSCs at different seeding densities on soft, thick hydrogels in basal media at 5 min and 20 hours. Coloured arrows indicate the increase in hydrogel deformations. Scale arrow=3. B) Quantification of displacements revealed that BMSCs created greater hydrogel deformations at lower seeding densities. Bars represent the mean and standard deviation of the 90^th^ percentiles of the hydrogel displacements. Significant differences between groups were determined using a 2-way ANOVA. ****=p<0.001.

### Cell-induced traction forces decrease with respect to time of culture

As osteogenic differentiation intrinsically relies on cell-cell contact and extended culture, we hypothesised that cell crowding during prolonged cell culture might abrogate cell mechanosensing by collective cell behaviour. To test this, we plated BMSCs at 5,000 cells/cm^2^ on thin and thick soft substrates in basal medium and tracked displacements every five minutes at 24 hrs, 10 days, and 7 weeks post-seeding. At early time points (24 hrs), hydrogel deformations were greater for cells on thick PAAm hydrogels than thin hydrogels (Figure 6A and B; also observed in Supplementary Video 4). As expected, cell density on all hydrogels increased after 10 days or 7 weeks compared to day 1, as observed in the representative pictures in Figure 6A. At ten days, there were still significantly greater displacements measured on thick compared to thin gels; however, by 7 weeks, there was no significant difference (p < 0.05). In parallel, there was a significant decrease in mean displacements for cells on thick hydrogels concerning time between day 10 and week 7 (Figure 6B; 19.8 ± 0.9 µm on day 1 compared to 5.2 ± 0.1 µm on week seven (thick hydrogels); 8.9 ± 0.8 µm on day 1 compared to 4.7 ± 0.3 µm on week seven (thin hydrogels); p<0.0001).

**Figure 6.**
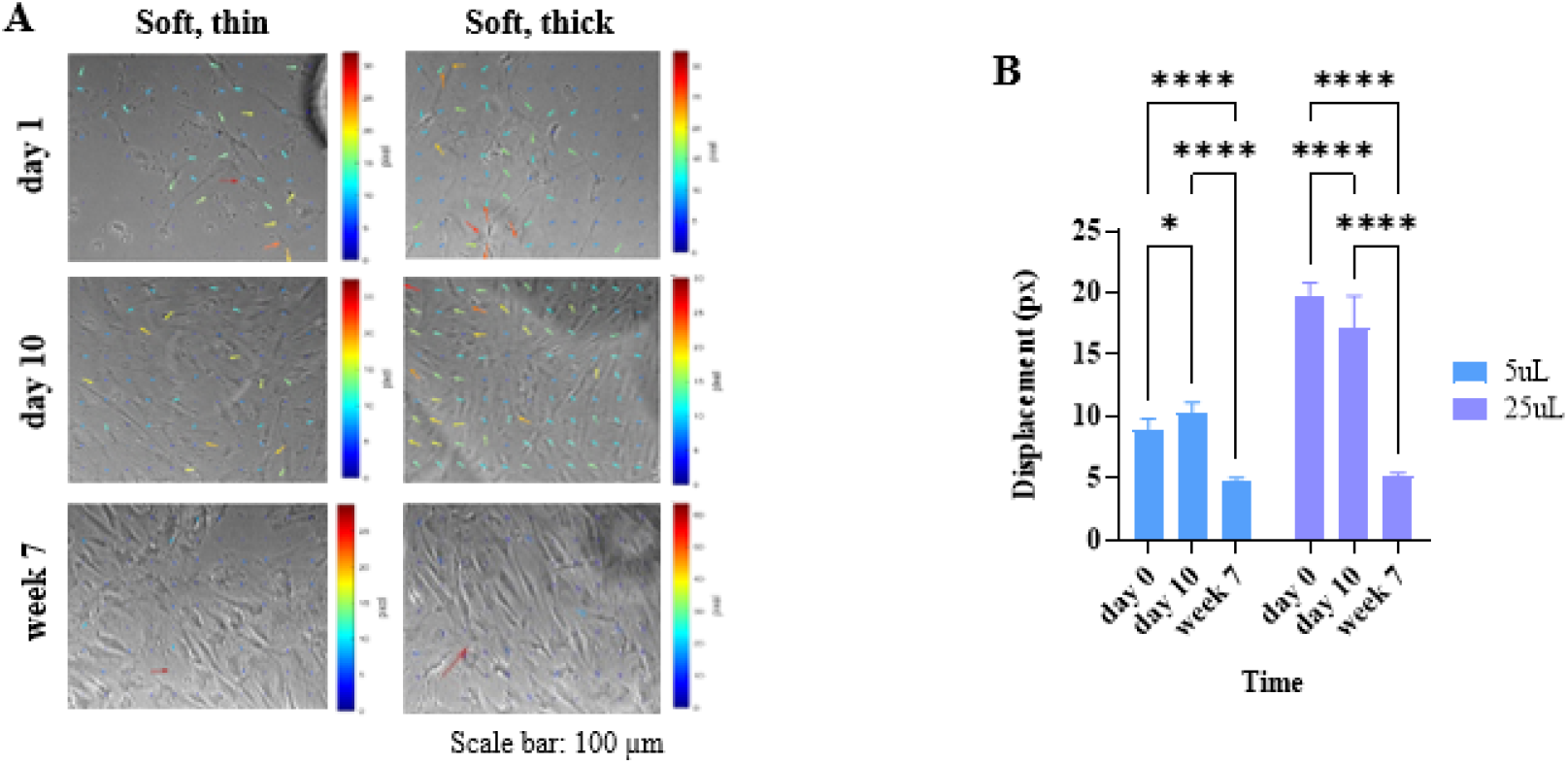
BMSC-induced hydrogel deformations decrease with respect to culture time. A) Time-lapse imaging of cells on soft, thin and thick hydrogels on day 1, day 10 and week 7 shows that cells increase in density during culture period and that the magnitude of deformations decreases. B) This is confirmed by quantification, where greater deformations are measured on soft, thick hydrogels compared to soft ones at day 1 and day 10. However these decline by week 7, where there is no significant difference between thick and thin hydrogels. Bars represent the mean and standard deviation of the 90^th^ percentiles of the hydrogel displacements on soft, thin and thick materials on day 1, day 10 and week 7.

We also compared time-dependent cell-induced displacements concerning time in basal and osteogenic medium. For low and moderate cell densities, there were significant increases in cell contractility in cells plated in the osteogenic medium after only 24 hours (Figure 7 A and B and Supplementary Video 5). As in other experiments, cell displacements’ magnitude declined over time, with a greater decline in cells plated at higher density. This effect was more pronounced in cells in osteogenic conditions, where displacements were significantly reduced compared to basal (3.6 ± 0.5 µm vs. 7.1 ± 0.6 µm).

**Figure 7.**
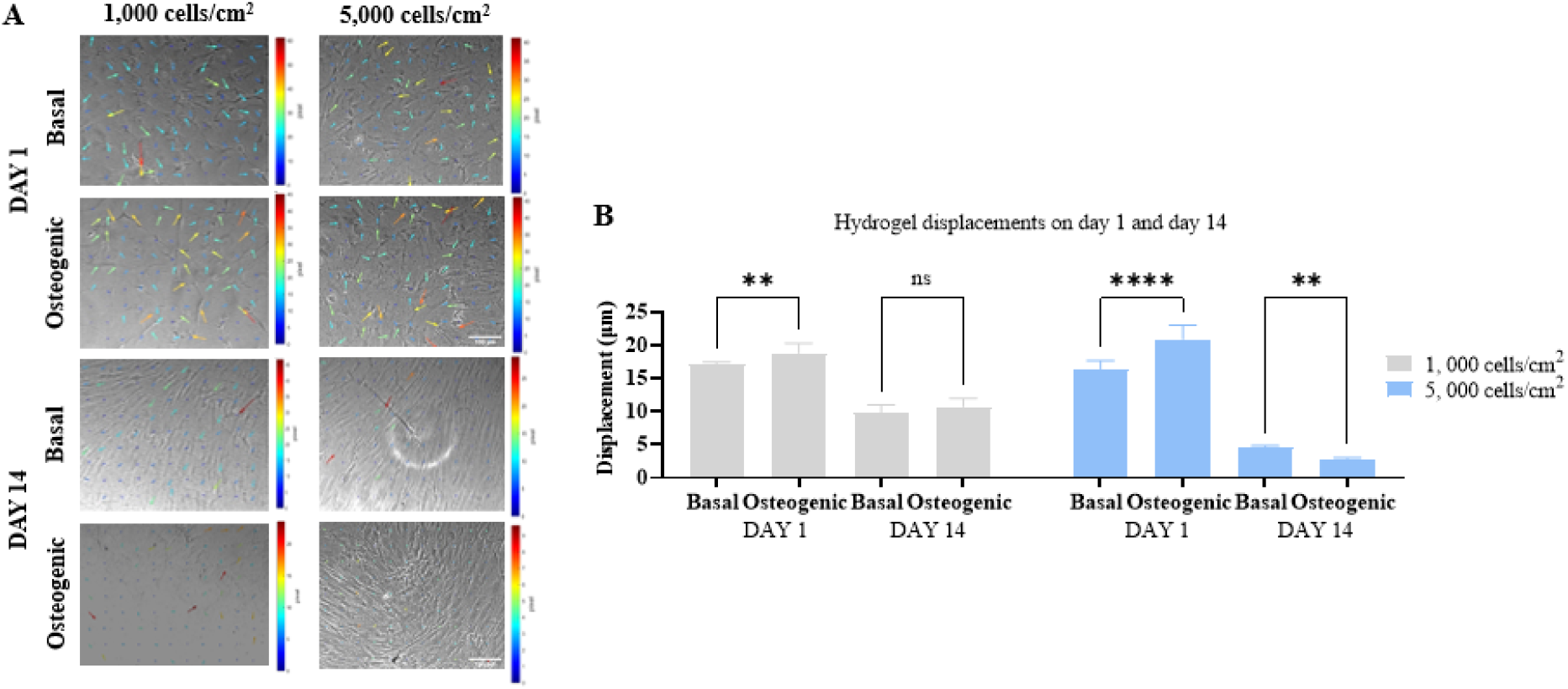
BMSCs generated different hydrogel deformations in basal and osteogenic conditions at different seeding densities. A) Time-lapse images of BMSCs on soft, thick PAAm hydrogels in basal and osteogenic media at low seeding densities (1,000 and 5,000 cells/cm^2^) on days 1 and 14. Cell morphology and alignment varied with the increase in seeding density. Coloured arrows show displacements on soft, thick hydrogels on day 1 but not on day 14. Images were obtained at 10X magnification with a Nikon Eclipse Ti inverted microscope. Scale arrow=3. B) Hydrogel displacements by BMSCs decreased on day 14 compared to day 1, regardless of the seeding density. Bars represent the mean and standard deviation of the 90^th^ percentiles of the hydrogel displacements on soft, thick materials on days 1 and day 14 in basal and osteogenic conditions.

This data indicates a time-dependent decrease in cell-induced hydrogel displacements for cells on thick materials, which is associated with an increase in cell density and the addition of osteogenic medium.

## DISCUSSION

The elastic modulus of extracellular matrix and biomaterials is known to be a factor in directing the differentiation of BMSCs, with materials of higher modulus promoting osteogenic differentiation. However, material geometry affects the true stiffness that individual and groups of cells ‘feel’. In this study, we found that matrix deformations exerted by BMSCs are constrained by both material thickness and cell density, which may provide a mechanism for why in some circumstances BMSCs become insensitive to substrate stiffness during differentiation to the osteoblastic lineage.

In order to achieve these conclusions, we first observed that soft hydrogels made from equivalent concentrations of monomer and crosslinker were thicker than their stiff counterparts. This is likely due to swelling. Softer hydrogels have been shown to have larger pores than stiff hydrogels, and hold more water molecules ^30^. Protick et al., 2022 showed that the swelling ratio decreased on stiffer hydrogels; ∼900% for soft hydrogels, and ∼350% for stiff hydrogels ^31^. In parallel with swelling, we also found that soft hydrogels exhibited wrinkles on their surfaces. This agrees with a previous report showing that during soft hydrogel fabrication, wrinkles appear to the differences in osmotic pressure in buffer or media solutions, causing water ingress and thus swelling ^32^. We noticed that the number of wrinkles decreased and their length increased as a function of increasing hydrogel thickness. As gels are covalently coupled to the underlying glass support, the hydrogel can only swell in a direction apical to the glass surface, resulting in a compressive stress in the hydrogel layer which may then induce buckling at the surface of the hydrogel. Regardless of these differences, we found previously that these hydrogels differ considerably in their stiffness ^28^, and exhibit large, flat areas easily distinguishable from folder areas, making them suitable for cell culture studies.

The observation that BMSCs respond to materials of different elastic modulus by modulating the degree of spreading and actin fibre generation is well understood ^33,34^. Cells on materials of a low elastic modulus are unable to generate cytoskeletal tension and generally appear smaller. In contrast, cells on materials with higher moduli generate higher forces that promote the formation of rigid, stiff, and contractile stress fibres, which promote cell spreading ^2,35^. In addition, our data showing that cells spread to a greater extent on thick, soft materials compared to thin ones, also reflects extensive literature supporting the ability of cells to detect boundaries ^8,9,36^. It has previously been assumed that this is likely due to constraint to cell-induced lateral hydrogel deformation imposed by an underlying stiff support. ^8^ Our data showing reduced displacements on materials of identical composition but different thicknesses, indicate that this is unexplored hypothesis is likely correct, and reflects our earlier data showing that groups of osteoscarcoma cells impose smaller displacements on thin compared to thick hydrogels.

Numerous previous studies reported that changes in ECM stiffness influence the differentiation potential of BMSCs (also sometimes known as mesenchymal stem cells (MSCs)) ^5,37^. Differentiation is subject to changes in the ECM stiffness to differentiate to the specific cell type that matches the tissue stiffness; 0.1-1 kPa hydrogels are neurogenic, 8-17 kPa are myogenic, and 25-40 kPa are osteogenic ^5^. We evaluated the osteogenic differentiation potential of BMSCs by quantifying ALP activity, a marker for BMSC osteogenic differentiation. Despite other data reporting that stiff hydrogels promote an increase in cell proliferation and osteogenic differentiation, we found no significant difference in ALP activity between soft and stiff hydrogels with different thicknesses. Previous reports indicate that high seeding density, may abrogate the ability of cells to detect soft materials ^27^.

These results led us to quantify hydrogel displacements to test whether cell-induced hydrogel displacements were inhibited with respect to cell density. We observed that cells created greater deformations on soft, thick hydrogel deformations at low seeding density compared to high seeding density in both basal and osteogenic medium. This likely reflects inhibition of cell contractility due to a ‘tug-of-war’ between neighbouring cells ^12^, with cells mechanically coupled both by direct contact or through the underlying material ^17^. We suspect that in subconfluent monolayers cells BMSCs may in fact be mechanically ‘coupled’ across the entirely of the hydrogel with the underlying glass support, detecting a higher stiffness than the independently-measured modulus of the material might suggest. Further experiments that may include measurements of whole-gel contraction, or control of the lateral dimensions of BMSCs layers, would be required to test this hypothesis formally.

In most experimental protocols, terminal osteogenic differentiation of BMSCs usually requires prolonged cell culture (2-3 weeks or more) and cell-cell contact. The observation that displacements also declined with respect to culture time (which is positively correlated with cell density) also supports this hypothesis. However, it is challenging here to rule out any effect of extracellular matrix deposition on the surface of the PAAm. It may be the case that at later timepoints, cells have secreted extracellular matrix which provides a stiffer growth substratum for the cells and which prevents direct mechanical coupling between cells and the underlying (fiduciary marker labelled) PAAm ^38^.

Further to these data, we found that BMSCs exerted significantly greater surface displacements when plated in medium containing osteogenic supplements than in basal medium. We consider that this is likely due to the glucocoticoid, dexamethasone, which has been shown to increase cell contractility in a range of cells, including mesenchymal stem cells ^39^ and alveolar epithelial cells ^40^, possibly through a role in modulating the formation and stability of F-actin ^41^. In addition, it was also evident that displacements were either unchanged or significantly lower for cells cultured in osteogenic medium compared to basal medium at later timepoints. It likely that this was due to increased proliferation in osteogenic conditions ^42^, leading to higher cell density and reduced measured displacements, by the mechanism we propose above.

In summary, cell mechanosensing is a complex process involving different variables such as ECM modulus and thickness, cell density and shape and the presence of supplements, determining cell traction forces and cell differentiation.

## ACKNOWLEDGEMENTS

We acknowledge CONAHCyT (National Council of humanities, Sciences and Technologies), Mexico and the Faculty of Medicine, Southampton University, UK, for funding. We also would like to thank Prof Ed Sander and Dr Hoda Zarkoob of the University of Iowa for advice with the MATLAB scripts used for quantifying displacements.

## REFERENCES

(1) Ringer, P.; Colo, G.; Fässler, R.; Grashoff, C. Sensing the Mechano-Chemical Properties of the Extracellular Matrix. Matrix Biol 2017, 64, 6–16. 10.1016/j.matbio.2017.03.004.

(2) Pelham, R. J.; Wang, Y. l. Cell Locomotion and Focal Adhesions Are Regulated by Substrate Flexibility. Proc Natl Acad Sci U S A 1997, 94 (25), 13661–13665. 10.1073/pnas.94.25.13661.

(3) Lo, C.-M.; Wang, H.-B.; Dembo, M.; Wang, Y. Cell Movement Is Guided by the Rigidity of the Substrate. Biophysical Journal 2000, 79 (1), 144–152. 10.1016/S0006-3495(00)76279-5.

(4) Wang, Y.; Wang, G.; Luo, X.; Qiu, J.; Tang, C. Substrate Stiffness Regulates the Proliferation, Migration, and Differentiation of Epidermal Cells. Burns 2012, 38 (3), 414–420. 10.1016/j.burns.2011.09.002.

(5) Engler, A. J.; Sen, S.; Sweeney, H. L.; Discher, D. E. Matrix Elasticity Directs Stem Cell Lineage Specification. Cell 2006, 126 (4), 677–689. 10.1016/j.cell.2006.06.044.

(6) Evans, N. D.; Minelli, C.; Gentleman, E.; LaPointe, V.; Patankar, S. N.; Kallivretaki, M.; Chen, X.; Roberts, C. J.; Stevens, M. M. Substrate Stiffness Affects Early Differentiation Events in Embryonic Stem Cells. Eur Cell Mater 2009, 18, 1–13; discussion 13-14. 10.22203/ecm.v018a01.

(7) Swift, J.; Ivanovska, I. L.; Buxboim, A.; Harada, T.; Dingal, P. C. D. P.; Pinter, J.; Pajerowski, J. D.; Spinler, K. R.; Shin, J.-W.; Tewari, M.; Rehfeldt, F.; Speicher, D. W.; Discher, D. E. Nuclear Lamin-A Scales with Tissue Stiffness and Enhances Matrix-Directed Differentiation. Science 2013, 341 (6149), 1240104. 10.1126/science.1240104.

(8) Leong, W. S.; Tay, C. Y.; Yu, H.; Li, A.; Wu, S. C.; Duc, D.-H.; Lim, C. T.; Tan, L. P. Thickness Sensing of hMSCs on Collagen Gel Directs Stem Cell Fate. Biochem Biophys Res Commun 2010, 401 (2), 287–292. 10.1016/j.bbrc.2010.09.052.

(9) Buxboim, A.; Rajagopal, K.; Brown, A. E. X.; Discher, D. E. How Deeply Cells Feel: Methods for Thin Gels. J. Phys.: Condens. Matter 2010, 22 (19), 194116. 10.1088/0953-8984/22/19/194116.

(10) Lin, Y.-C.; Tambe, D. T.; Park, C. Y.; Wasserman, M. R.; Trepat, X.; Krishnan, R.; Lenormand, G.; Fredberg, J. J.; Butler, J. P. Mechanosensing of Substrate Thickness. Phys Rev E Stat Nonlin Soft Matter Phys 2010, 82 (4 0 1), 041918.

(11) Evans, N. D.; Gentleman, E. The Role of Material Structure and Mechanical Properties in Cell-Matrix Interactions. J Mater Chem B 2014, 2 (17), 2345–2356. 10.1039/c3tb21604g.

(12) Trepat, X.; Wasserman, M. R.; Angelini, T. E.; Millet, E.; Weitz, D. A.; Butler, J. P.; Fredberg, J. J. Physical Forces during Collective Cell Migration. Nature Phys 2009, 5 (6), 426–430. 10.1038/nphys1269.

(13) Tusan, C. G.; Man, Y.-H.; Zarkoob, H.; Johnston, D. A.; Andriotis, O. G.; Thurner, P. J.; Yang, S.; Sander, E. A.; Gentleman, E.; Sengers, B. G.; Evans, N. D. Collective Cell Behavior in Mechanosensing of Substrate Thickness. Biophys J 2018, 114 (11), 2743–2755. 10.1016/j.bpj.2018.03.037.

(14) Zarkoob, H.; Bodduluri, S.; Ponnaluri, S. V.; Selby, J. C.; Sander, E. A. Substrate Stiffness Affects Human Keratinocyte Colony Formation. Cell Mol Bioeng 2015, 8 (1), 32–50. 10.1007/s12195-015-0377-8.

(15) Mertz, A. F.; Banerjee, S.; Che, Y.; German, G. K.; Xu, Y.; Hyland, C.; Marchetti, M. C.; Horsley, V.; Dufresne, E. R. Scaling of Traction Forces with the Size of Cohesive Cell Colonies. Phys. Rev. Lett. 2012, 108 (19), 198101. 10.1103/PhysRevLett.108.198101.

(16) Emerman, J. T.; Pitelka, D. R. Maintenance and Induction of Morphological Differentiation in Dissociated Mammary Epithelium on Floating Collagen Membranes. In Vitro 1977, 13 (5), 316–328. 10.1007/BF02616178.

(17) Reinhart-King, C. A.; Dembo, M.; Hammer, D. A. Cell-Cell Mechanical Communication through Compliant Substrates. Biophys J 2008, 95 (12), 6044–6051. 10.1529/biophysj.107.127662.

(18) El-Rashidy, A. A.; El Moshy, S.; Radwan, I. A.; Rady, D.; Abbass, M. M. S.; Dörfer, C. E.; Fawzy El-Sayed, K. M. Effect of Polymeric Matrix Stiffness on Osteogenic Differentiation of Mesenchymal Stem/Progenitor Cells: Concise Review. Polymers 2021, 13 (17), 2950. 10.3390/polym13172950.

(19) Wen, J. H.; Vincent, L. G.; Fuhrmann, A.; Choi, Y. S.; Hribar, K. C.; Taylor-Weiner, H.; Chen, S.; Engler, A. J. Interplay of Matrix Stiffness and Protein Tethering in Stem Cell Differentiation. Nature Mater 2014, 13 (10), 979–987. 10.1038/nmat4051.

(20) Lee, J.; Abdeen, A. A.; Huang, T. H.; Kilian, K. A. Controlling Cell Geometry on Substrates of Variable Stiffness Can Tune the Degree of Osteogenesis in Human Mesenchymal Stem Cells. J Mech Behav Biomed Mater 2014, 38, 209–218. 10.1016/j.jmbbm.2014.01.009.

(21) Witkowska-Zimny, M.; Walenko, K.; Wrobel, E.; Mrowka, P.; Mikulska, A.; Przybylski, J. Effect of Substrate Stiffness on the Osteogenic Differentiation of Bone Marrow Stem Cells and Bone-Derived Cells. Cell Biology International 2013, 37 (6), 608–616. 10.1002/cbin.10078.

(22) Rowlands, A. S.; George, P. A.; Cooper-White, J. J. Directing Osteogenic and Myogenic Differentiation of MSCs: Interplay of Stiffness and Adhesive Ligand Presentation. American Journal of Physiology-Cell Physiology 2008, 295 (4), C1037–C1044. 10.1152/ajpcell.67.2008.

(23) Barreto, S.; Gonzalez-Vazquez, A.; Cameron, A. R.; Cavanagh, B.; Murray, D. J.; O’Brien, F. J. Identification of the Mechanisms by Which Age Alters the Mechanosensitivity of Mesenchymal Stromal Cells on Substrates of Differing Stiffness: Implications for Osteogenesis and Angiogenesis. Acta Biomater 2017, 53, 59–69. 10.1016/j.actbio.2017.02.031.

(24) Viale-Bouroncle, S.; Völlner, F.; Möhl, C.; Küpper, K.; Brockhoff, G.; Reichert, T. E.; Schmalz, G.; Morsczeck, C. Soft Matrix Supports Osteogenic Differentiation of Human Dental Follicle Cells. Biochem Biophys Res Commun 2011, 410 (3), 587–592. 10.1016/j.bbrc.2011.06.031.

(25) Stanton, A. E.; Tong, X.; Yang, F. Extracellular Matrix Type Modulates Mechanotransduction of Stem Cells. Acta Biomater 2019, 96, 310–320. 10.1016/j.actbio.2019.06.048.

(26) Mao, A. S.; Shin, J.-W.; Mooney, D. J. Effects of Substrate Stiffness and Cell-Cell Contact on Mesenchymal Stem Cell Differentiation. Biomaterials 2016, 98, 184–191. 10.1016/j.biomaterials.2016.05.004.

(27) Venugopal, B.; Mogha, P.; Dhawan, J.; Majumder, A. Cell Density Overrides the Effect of Substrate Stiffness on Human Mesenchymal Stem Cells’ Morphology and Proliferation. Biomater. Sci. 2018, 6 (5), 1109–1119. 10.1039/C7BM00853H.

(28) Xu, D.; Hernandez Miranda, M.; Evans, N. D.; Sengers, B. G.; Browne, M.; Cook, R. B. Depth Profiling via Nanoindentation for Characterisation of the Elastic Modulus and Hydraulic Properties of Thin Hydrogel Layers. Journal of the Mechanical Behavior of Biomedical Materials Under consideration.

(29) Janeczek, A. A.; Tare, R. S.; Scarpa, E.; Moreno-Jimenez, I.; Rowland, C. A.; Jenner, D.; Newman, T. A.; Oreffo, R. O. C.; Evans, N. D. Transient Canonical Wnt Stimulation Enriches Human Bone Marrow Mononuclear Cell Isolates for Osteoprogenitors. Stem Cells 2016, 34 (2), 418–430. 10.1002/stem.2241.

(30) Subramani, R.; Izquierdo-Alvarez, A.; Bhattacharya, P.; Meerts, M.; Moldenaers, P.; Ramon, H.; Van Oosterwyck, H. The Influence of Swelling on Elastic Properties of Polyacrylamide Hydrogels. Frontiers in Materials 2020, 7.

(31) Protick, F. K.; Amit, S. K.; Amar, K.; Nath, S. D.; Akand, R.; Davis, V. A.; Nilufar, S.; Chowdhury, F. Additive Manufacturing of Viscoelastic Polyacrylamide Substrates for Mechanosensing Studies. ACS Omega 2022, 7 (28), 24384–24395. 10.1021/acsomega.2c01817.

(32) Saha, K.; Kim, J.; Irwin, E.; Yoon, J.; Momin, F.; Trujillo, V.; Schaffer, D. V.; Healy, K. E.; Hayward, R. C. Surface Creasing Instability of Soft Polyacrylamide Cell Culture Substrates. Biophys J 2010, 99 (12), L94–96. 10.1016/j.bpj.2010.09.045.

(33) Wang, H.-B.; Dembo, M.; Wang, Y.-L. Substrate Flexibility Regulates Growth and Apoptosis of Normal but Not Transformed Cells. American Journal of Physiology - Cell Physiology 2000, 279 (5 48-5), C1345–C1350. 10.1152/ajpcell.2000.279.5.c1345.

(34) Yeung, T.; Georges, P. C.; Flanagan, L. A.; Marg, B.; Ortiz, M.; Funaki, M.; Zahir, N.; Ming, W.; Weaver, V.; Janmey, P. A. Effects of Substrate Stiffness on Cell Morphology, Cytoskeletal Structure, and Adhesion. Cell Motility 2005, 60 (1), 24–34. 10.1002/cm.20041.

(35) Zhou, D. W.; Lee, T. T.; Weng, S.; Fu, J.; García, A. J. Effects of Substrate Stiffness and Actomyosin Contractility on Coupling between Force Transmission and Vinculin-Paxillin Recruitment at Single Focal Adhesions. Mol Biol Cell 2017, 28 (14), 1901–1911. 10.1091/mbc.E17-02-0116.

(36) Merkel, R.; Kirchgeßner, N.; Cesa, C. M.; Hoffmann, B. Cell Force Microscopy on Elastic Layers of Finite Thickness. Biophysical Journal 2007, 93 (9), 3314–3323. 10.1529/biophysj.107.111328.

(37) Lanniel, M.; Huq, E.; Allen, S.; Buttery, L.; M. Williams, P.; R. Alexander, M. Substrate Induced Differentiation of Human Mesenchymal Stem Cells on Hydrogels with Modified Surface Chemistry and Controlled Modulus. Soft Matter 2011, 7 (14), 6501–6514. 10.1039/C1SM05167A.

(38) Li, B.; Moshfegh, C.; Lin, Z.; Albuschies, J.; Vogel, V. Mesenchymal Stem Cells Exploit Extracellular Matrix as Mechanotransducer. Sci Rep 2013, 3 (1), 2425. 10.1038/srep02425.

(39) Sridharan Weaver, S.; Li, Y.; Foucard, L.; Majeed, H.; Bhaduri, B.; Levine, A. J.; Kilian, K. A.; Popescu, G. Simultaneous Cell Traction and Growth Measurements Using Light. J Biophotonics 2019, 12 (3), e201800182. 10.1002/jbio.201800182.

(40) Puig, F.; Gavara, N.; Sunyer, R.; Carreras, A.; Farré, R.; Navajas, D. Stiffening and Contraction Induced by Dexamethasone in Alveolar Epithelial Cells. Exp Mech 2009, 49 (1), 47–55. 10.1007/s11340-007-9072-6.

(41) Castellino, F.; Heuser, J.; Marchetti, S.; Bruno, B.; Luini, A. Glucocorticoid Stabilization of Actin Filaments: A Possible Mechanism for Inhibition of Corticotropin Release. Proc Natl Acad Sci U S A 1992, 89 (9), 3775–3779. 10.1073/pnas.89.9.3775.

(42) Nishimura, I.; Hisanaga, R.; Sato, T.; Arano, T.; Nomoto, S.; Ikada, Y.; Yoshinari, M. Effect of Osteogenic Differentiation Medium on Proliferation and Differentiation of Human Mesenchymal Stem Cells in Three-Dimensional Culture with Radial Flow Bioreactor. Regen Ther 2015, 2, 24–31. 10.1016/j.reth.2015.09.001.

