## Supplemental Figures for "Geometric constraint of mechanosensing in bone marrow stromal cell cultures prevents stiffness-induced differentiation"

### SUPPLEMENTARY FIGURE 1

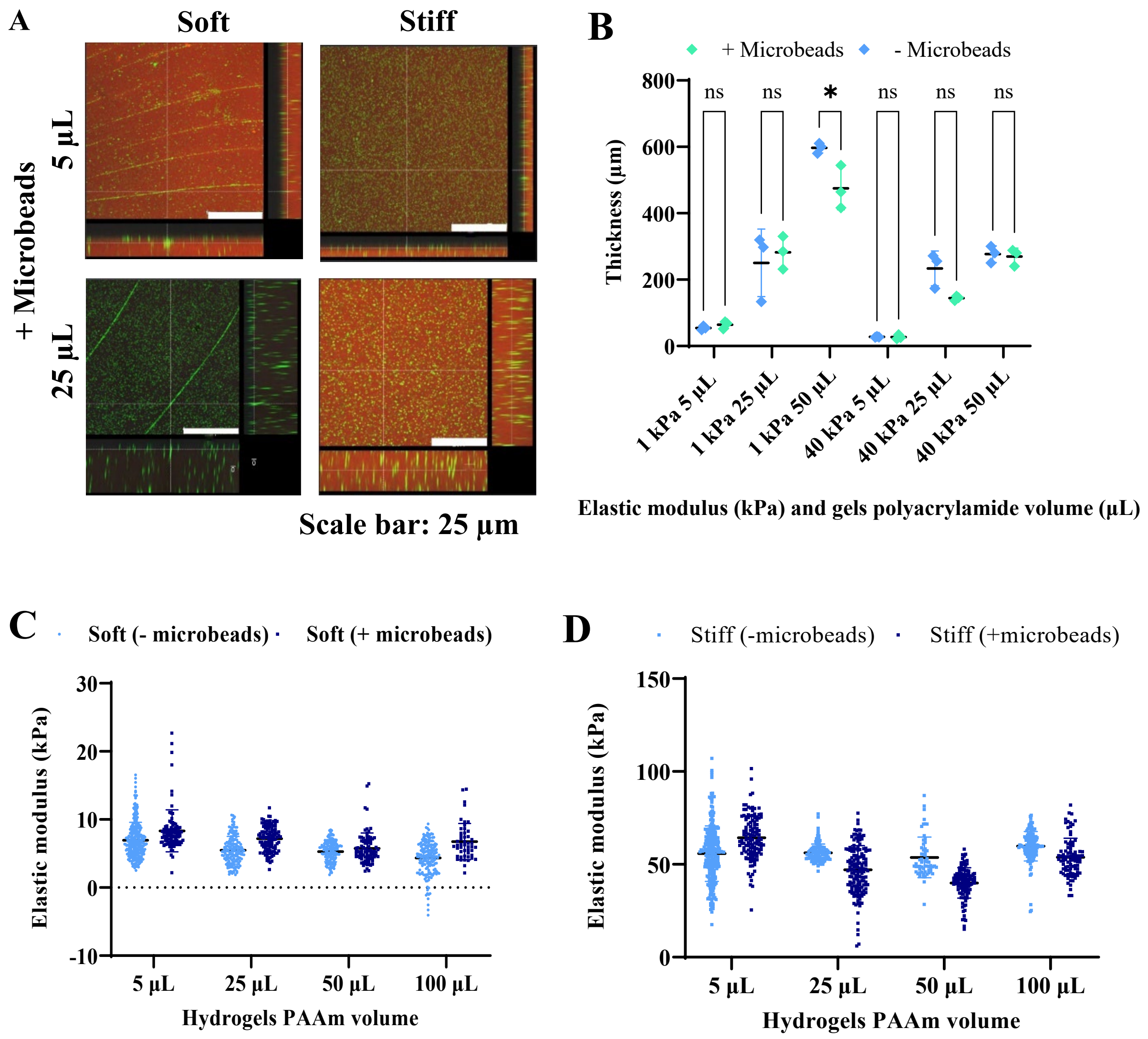


***Supplementary Figure 1.*** *Fluorescent microbeads are distributed evenly in stiff hydrogels but show alignment with gel wrinkles in soft hydrogels (A). Microbead incorporation did not affect the thickness of either thin or thick hydrogels, except for in 50 uL soft hydrogels (B). Microbead incorporation had little effect on corrected elastic modulus on either soft (C) or thick (D) hydrogels. Statistical differences were calculated by the 2-way ANOVA method (****=p<0.0001).*

### SUPPLEMENTARY FIGURE 2

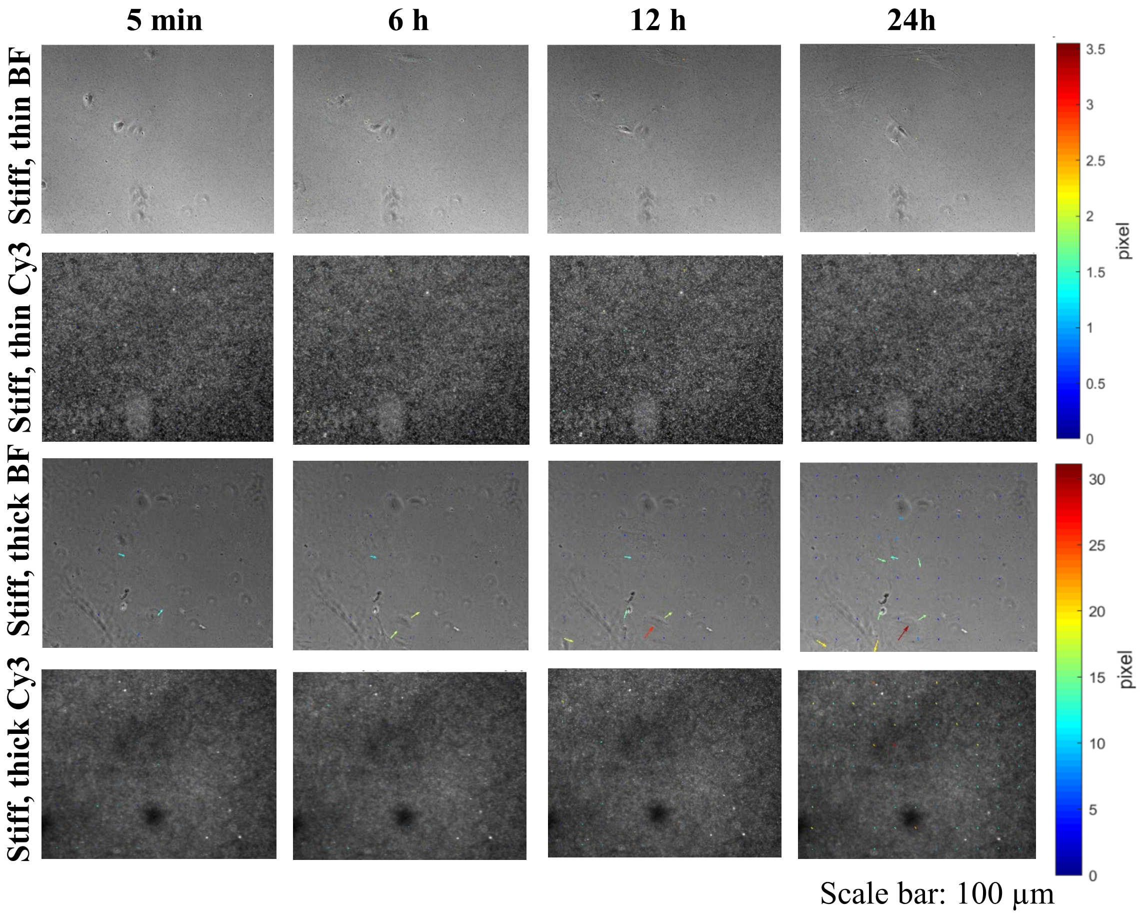


***Supplementary Figure 2.*** *Stiff matrices exhibit small deformations by BMSCs regardless of hydrogel thickness. Small coloured arrows show small deformations on stiff, thick compared to the thin counterparts hydrogels. Phase contrast (BF) and fluorescent (Cy3) images were obtained at 10X magnification under a Nikon Eclipse Ti inverted microscope. Scale bar=3.*
